# Coloring inside the lines: genomic architecture and evolution of a widespread color pattern in frogs

**DOI:** 10.1101/2021.10.28.466315

**Authors:** Sandra Goutte, Imtiyaz Hariyani, Kole Deroy Utzinger, Yann Bourgeois, Stéphane Boissinot

## Abstract

Traits shared among distantly related lineages are indicators of common evolutionary constraints, at the ecological, physiological or molecular level. The vertebral stripe is a color pattern that is widespread across the anuran phylogeny. Despite its prevalence in the order, surprisingly little is known about the genetic basis and evolutionary dynamic of this color pattern. Here we combine histology, genome- and transcriptome-wide analyses with order-scale phylogenetic comparative analyses to investigate this common phenotype. We show that the vertebral stripe has evolved hundreds of times in the evolutionary history of anurans and is selected for in terrestrial habitats. Using the Ethiopian *Ptychadena* radiation as a model system, we demonstrate that variation at the *ASIP* gene is responsible for the different vertebral stripe phenotypes. Alleles associated to these phenotypes are younger than the split between closely related *Ptychadena* species, thus indicating that the vertebral stripe results from parallel evolution within the group. Our findings demonstrate that this widespread color pattern evolves rapidly and recurrently in terrestrial anurans, and therefore constitute an ideal system to study repeated evolution.

## Introduction

Animal color patterns are conspicuous hallmarks of selection. Color patterns may evolve because they are linked to a beneficial physiological trait^1, 2^ or because they serve as sexual^3, 4^ or warning signals^5^. Alternatively, color pattern can help avoid detection from visually-oriented predators or prey by disrupting body shape recognition^6–9^, masquerading as an object or animal^10^, countershading^11^, or substrate-matching^12–14^. In many species, multiple color patterns coexist within or between populations. These polymorphisms can be maintained by divergent mating strategies^15, 16^, apostatic selection (preference for the most common morph by predators), temporal or spatial habitat heterogeneity^5^, or heterozygote advantage on correlated traits^2^. This diversity of selective regimes makes color patterns an ideal system to investigate the evolutionary mechanisms underlying phenotypic evolution.

The vertebral stripe is a common color pattern found across numerous distantly related anuran amphibians around the world (Fig. 1a). Phenotypes shared across highly divergent lineages can result from ancestral alleles conserved over millions of years of evolution, or have evolved independently multiple times, perhaps driven by similar selective forces. While predator-mediated selection is a widely assumed mechanism for the evolution and maintenance of most color patterns in anurans^17^, the link between the anuran vertebral stripe and survival is only empirically supported in a few species^13, 18, 19^ (but see^20^). In order to understand the evolutionary history of the vertebral stripe in anurans, a broad-scale comparative analysis is necessary. Understanding the genomic architecture underlying phenotypes can also inform on the evolutionary mechanisms at play. Despite the commonness of the pattern, the genetic basis of the anuran vertebral stripe is largely unknown. In the few species investigated, the stripe is determined by a dominant allele at a single locus^17, 21^. However, as these inferences were made solely on crossings, the identity of the locus or loci involved remains to be determined.

**Figure 1.**
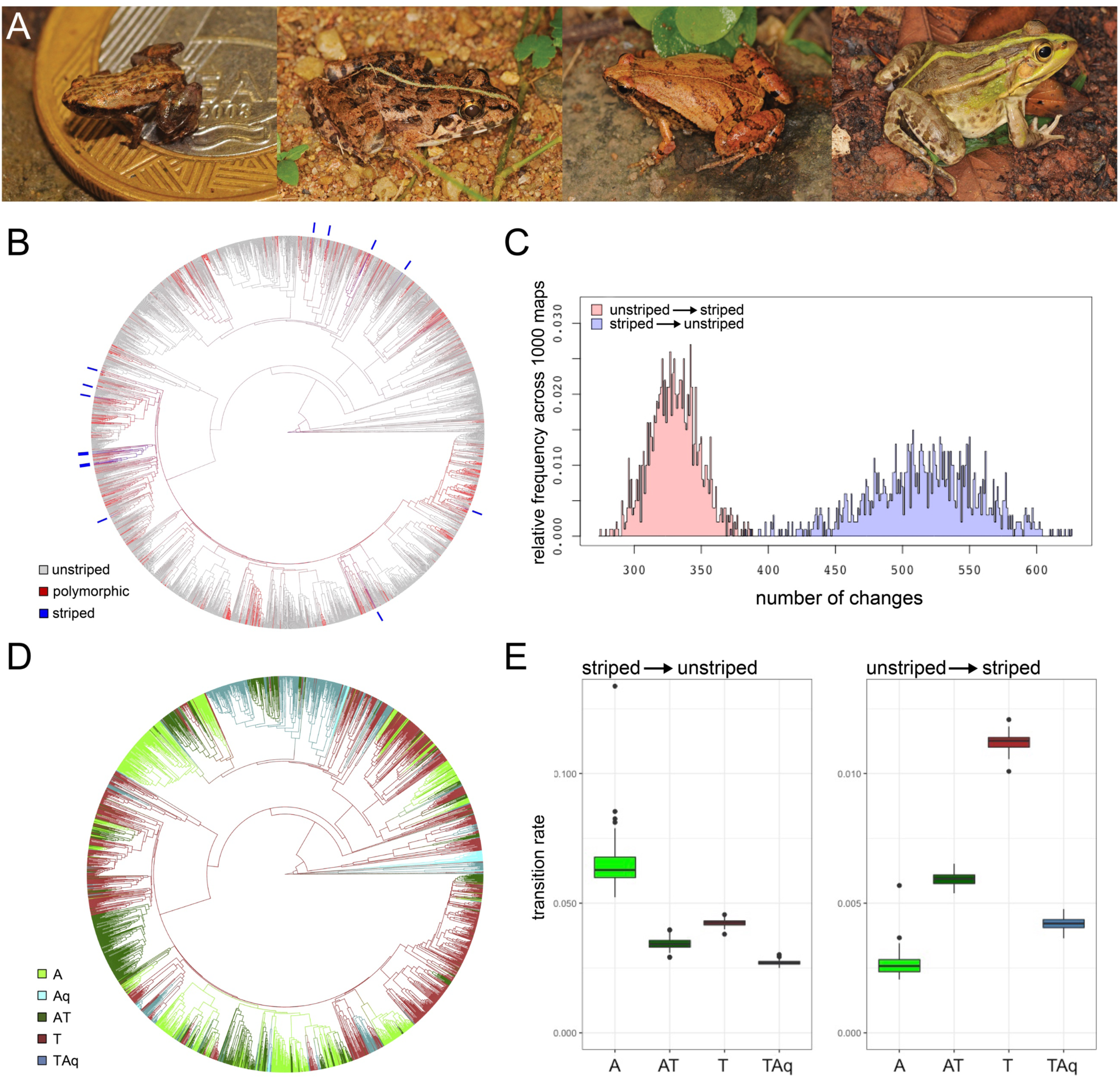
The evolution of the vertebral stripe in anurans. **A** Examples of the vertebral stripe in distantly related species: from left to right, *Brachycephalus hermogenesi* (family: Brachycephalidae), *Fejervarya limnocharis* (family: Dicroglossidae), *Microhyla ornata* (family: Microhylidae), *Pelophylax nigromaculatus* (family: Ranidae), **B** vertebral stripe morphs (unstriped = grey, striped = blue, polymorphic = red; monomorphic striped taxa are further indicated by blue bars for readability) mapped on the phylogeny of Anura (n=2,785 species; 1,000 stochastic maps), **C** estimated number of transitions between striped and unstriped morphs in the evolution of anurans based on 1,000 stochastic maps, **D** habitat use mapped on the phylogeny of Anura (n= 2,620 species; 100 stochastic maps), **E** transition rates between striped and unstriped phenotypes for each habitat, estimated for 100 stochastic maps. Habitat categories: A = arboreal, Aq = aquatic, AT = arboreal-terrestrial, T = terrestrial, TAq = terrestrial-aquatic.

Here we combine macro- and microevolutionary analyses with transcriptomic and histological data to investigate the evolutionary history and genomic architecture underlying the vertebral stripe in anurans. By integrating results at three different scales (order, species group, and species), our study exemplifies how natural selection combined with rapidly evolving genomic regions may result in recurrent phenotypic evolution.

## Results

### Evolutionary history of the vertebral stripe in anurans

To retrace the evolutionary history of the vertebral stripe in anurans, we examined the dorsal color pattern of 2,785 anuran species for which phylogenetic data was available^22^, representing 37.6% of species and 96.5% of families currently recognized in Anura^23^. A vertebral stripe morph was present in 15.7% of the 2,785 species included, and of those, 78% were polymorphic for the trait (Fig. 1b). Our analysis estimated that the vertebral stripe pattern evolved independently 330 ± 18 times across the phylogeny and was lost 515 ± 42 times (Fig. 1c). This result strongly supports the hypothesis of multiple origins of the anuran dorsal stripe and rejects the hypothesis of ancestral alleles conserved across anurans’ evolutionary history.

Once we established that the vertebral stripe evolved independently multiple times, we investigated the role of the environment in the evolution of this trait. We hypothesized that recurrent evolution of the vertebral stripe across anurans may be due to similar selective pressures in shared habitat types. To test this hypothesis, we carried a comparative analysis on 2,620 anuran species, and compared habitat-dependent and habitat-independent models of evolution for the vertebral stripe. Our analysis revealed that the rate of transition between unstriped and striped morphs is correlated with habitat (Supp. Fig. S1). The vertebral stripe evolved significantly more often in terrestrial clades compared to terrestrial-aquatic, arboreal, and terrestrial-arboreal linages (Fig 1d and 1e; Table 1). Arboreal lineages also showed the lowest gain and highest loss rates for the color pattern. The vertebral stripe may thus be selected against in arboreal habitats and selected for in terrestrial habitats.

**Table 1.**
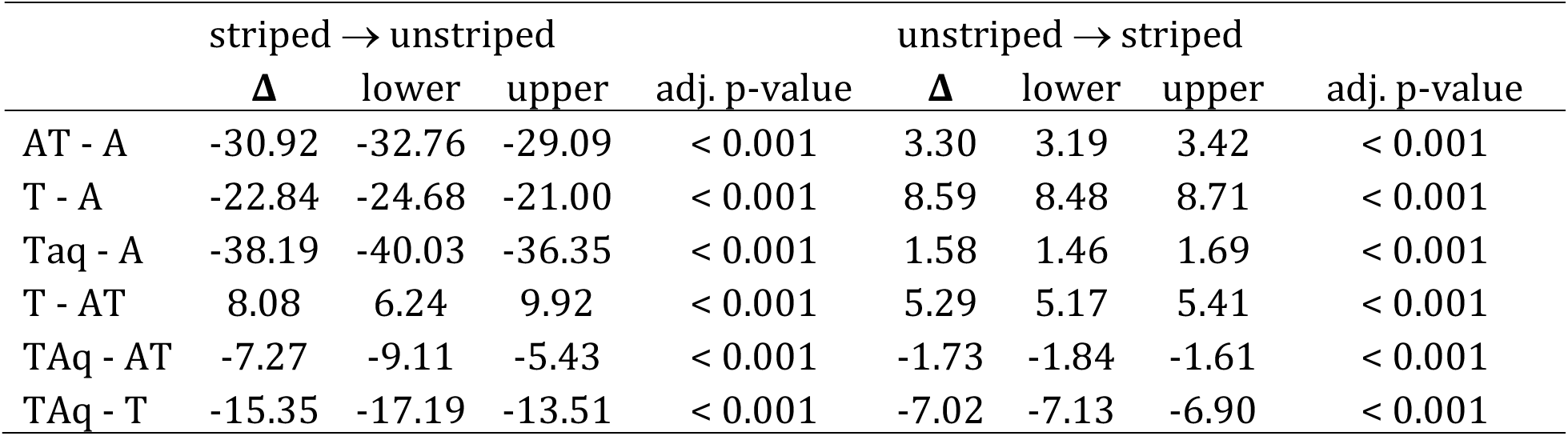
Tukey honest significant differences test of morph transition rates between habitat pairs. Values were multiplied by 10^3^ for readability. Average (Δ), lower and upper values of the difference between rates based on 100 stochastic maps are given. Habitat categories: A = arboreal, Aq = aquatic, AT = arboreal-terrestrial, T = terrestrial, TAq = terrestrial-aquatic.

### Cellular organization of the vertebral stripe

To investigate the molecular and cellular mechanisms underlying the vertebral stripe in anurans, we focused on a radiation of terrestrial frogs, the *Ptychadena neumanni* species complex. This monophyletic group encompasses 12 species^24^, which all present vertebral stripe polymorphism (the stripe could be thin, wide, or absent), except for two species: *P. harenna*, in which the vertebral stripe morph is always absent and *P. cooperi*, for which all individuals present a thin vertebral stripe (Fig. 2a and 2b). We examined the pigment cells organization in the dorsal skin of ten individuals of the *P. neumanni* species complex (*P. robeensis, P. nana, P. erlangeri* and *P. amharensis*) presenting different vertebral stripe phenotypes (thin or wide striped, or unstriped; Fig. 2b). Outside the stripe, numerous melanophores with dispersed melanosomes covering other chromatophores, and melanosomes in the epidermal layer create a dark coloration (Fig. 2c). Within the stripe, melanophores with aggregated melanosomes (when present) and no or few epidermal melanosomes result in a lighter shade (Fig. 2c). The number and state (aggregated vs. dispersed) of the melanophores, as well as the concentration of epidermal melanosomes thus seem to be the major determinants of the vertebral stripe pattern in the species examined.

**Figure 2.**
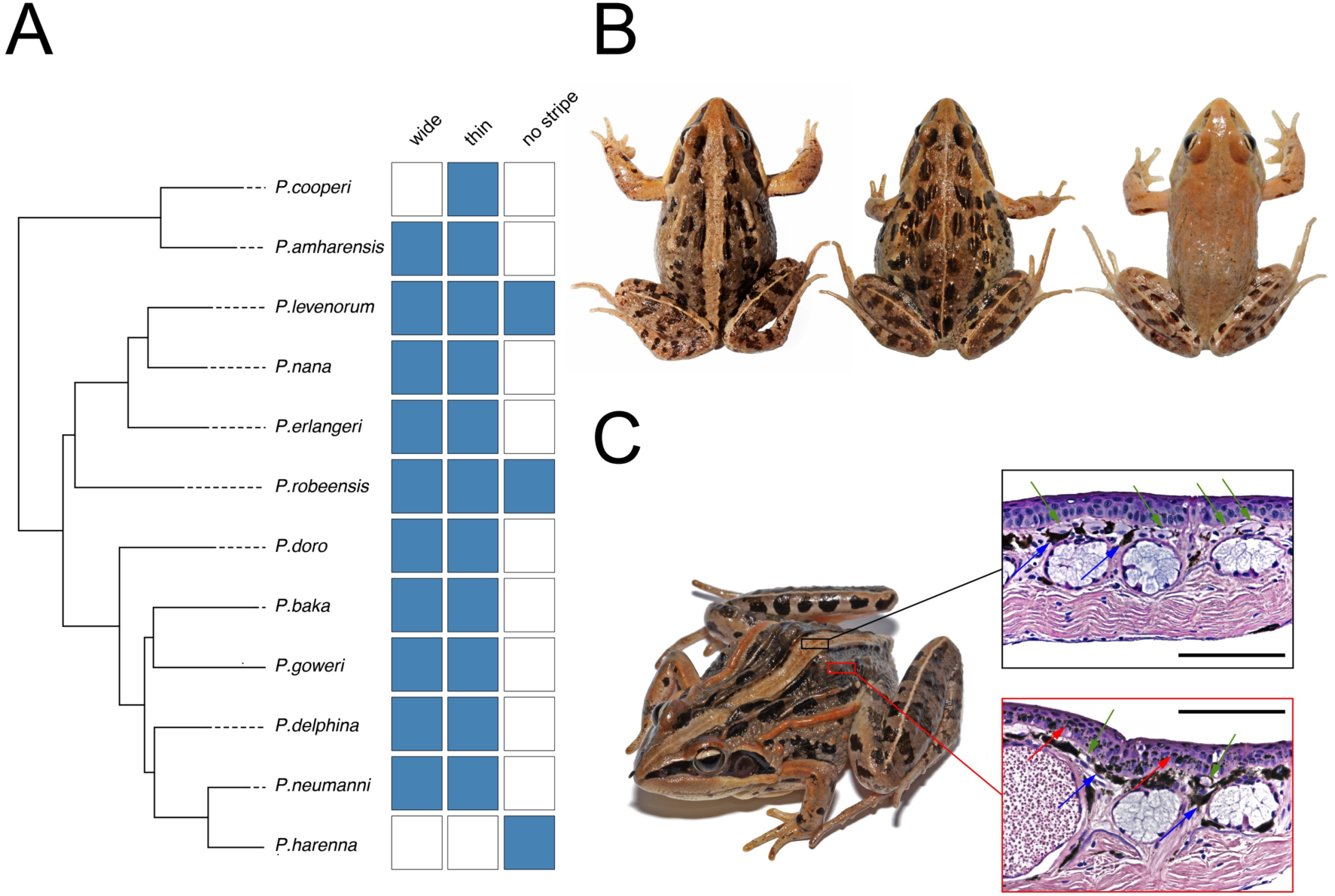
The vertebral stripe in Ethiopian *Ptychadena.* **A** Polymorphism of the vertebral stripe (wide or thin striped, or unstriped) in the *Ptychadena neumanni* species complex (phylogeny based on 500,000 genome-wide distributed SNPs, reproduced from^51^). Presence of the morph is indicated in blue, absence in white. **B** Adult *Ptychadena robeensis* presenting the three possible vertebral stripe morphs. From left to right: wide striped, thin striped, unstriped. **C** Histological sections of the dorsal skin within (top) and outside (bottom) the vertebral stripe in a female *Ptychadena erlangeri* (SB548; live photograph on the left). Scale bar = 200µm. Within the stripe (top), the few melanophores (blue arrows) are in a contracted state and do not entirely cover the xanthophores (green arrows), in contrast with the skin outside the stripe (bottom). Outside the stripe, numerous melanosomes (red arrows) are also present in the epidermal layer, creating a very dark coloration.

### Genomic architecture of the vertebral stripe in Ptychadena robeensis

To identify the genomic region(s) involved in the dorsal stripe pattern, we conducted a genome-wide association study (GWAS) on one species of the *P. neumanni* complex, *Ptychadena robeensis,* which is polymorphic for the trait. We produced whole-genome resequencing data (4.84X average coverage) for 52 individuals with either a wide (n = 25) or a thin (n = 27) vertebral stripe and aligned the reads on the recently assembled chromosome-level genome of the species^25^, resulting in a total of a 17,797,568 single-nucleotide polymorphisms (SNPs) dataset. The number of unstriped individuals collected being low (n=5), we excluded this phenotype from the analysis. We identified a single genomic region associated with the color pattern, which included multiple significant SNPs (Fig. 3a).

**Figure 3.**
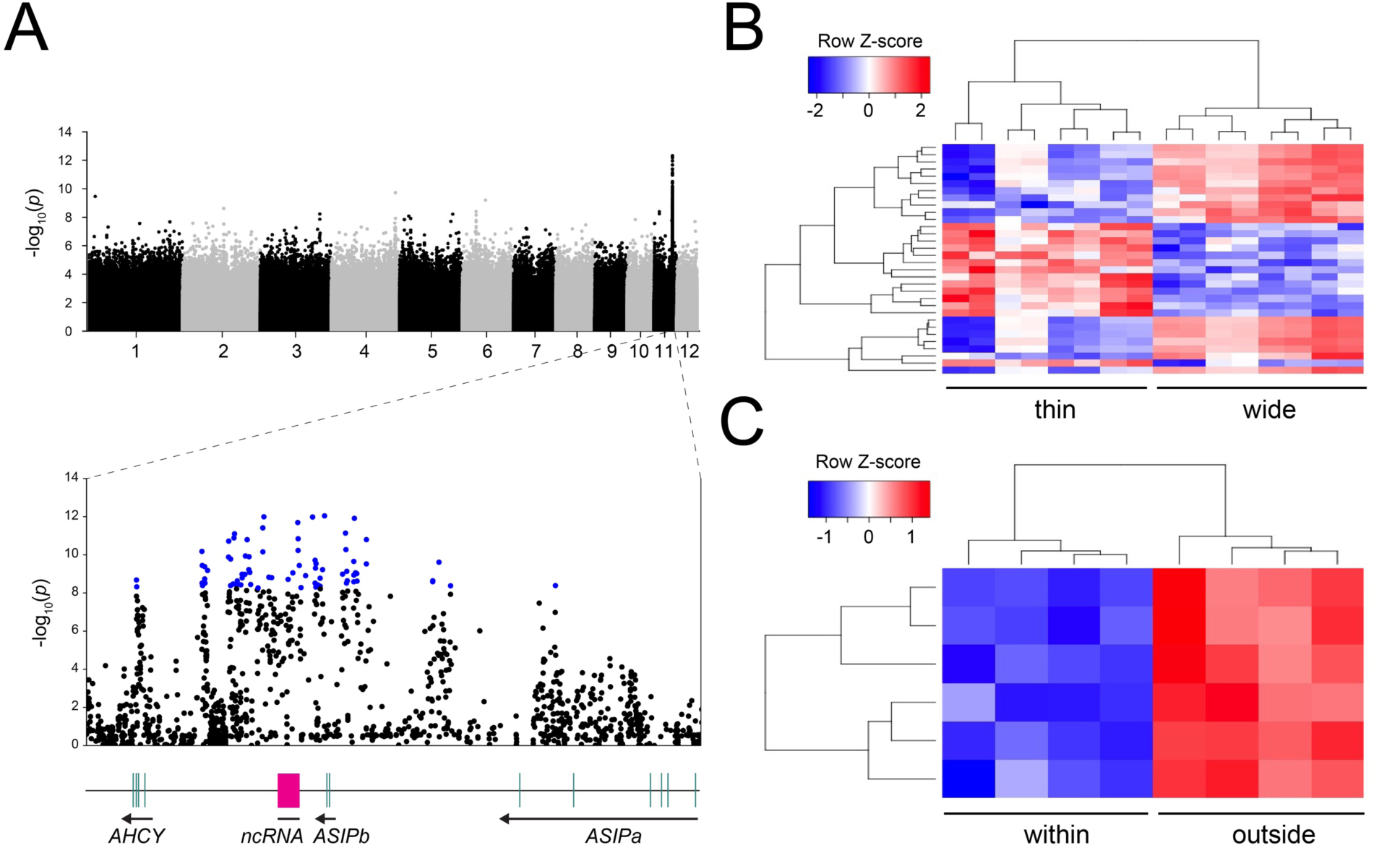
Genes associated with the stripe phenotype in *Ptychadena robeensis.* **A** Genome-wide association study (GWAS) reveals a single locus governing the vertebral stripe phenotype on chromosome 11 of *Ptychadena robeensis* (top panel). Significant SNPs (indicated in blue in the bottom panel) are located in between and downstream *ASIP* exons (indicated in light green below) and a non-coding RNA (pink). **B** and **C** Differential gene expression analysis in the skin of adult *Ptychadena robeensis.* 32 genes are differentially expressed between vertebral stripe phenotypes (**B**) and six within versus outside the vertebral stripe in the same individuals (**C**).

The identified SNPs are located on chromosome 11 downstream of a region containing two predicted^25^ copies of the *Agouti signaling protein* gene (*ASIPa* and *ASIPb*; see Methods and Supp. Fig. S3). The majority of the SNPs are in the region overlapping with *ASIPb* and a non-coding RNA (ncRNA) sequence (Fig. 3a) located downstream of this gene, and all of them fall outside coding sequences. *ASIP* is known to be involved in melanophore differentiation and melanin production in vertebrates^26, 27^. By examining the phenotypes of homozygote and heterozygote individuals, we determined that the *wide* allele is dominant over the *thin* allele. Because all of these SNPs are located outside the predicted protein coding regions, we hypothesize that they affect the expression of the gene. An up-regulation of *ASIP* is linked to lighter phenotypes in mammals^28, 29^ and fishes^27^, we could thus expect an up-regulation of *ASIP* in individuals with a wide vertebral stripe.

### Differential gene expression associated with the vertebral stripe

We explored transcriptome-wide patterns of gene expression in the skin of adult *P. robeensis* presenting a thin, wide, or no vertebral stripe (Fig. 3b and 3c). Surprisingly, the expression levels of *ASIPa*, *ASIPb* and the ncRNA were very low in dorsal skin, and no significant differential expression could be detected between phenotypes or between skin samples collected within versus outside the stripe (Supp. Fig. S4). The transcripts were at considerably greater concentration in the ventral skin, which is white and lacks any melanization (Supp. Fig. S4a). To quantify more precisely *ASIP* expression in the dorsal skin, we conducted a quantitative real-time PCR experiment (qPCR), which confirmed comparable expression levels of *ASIPb* within and outside the vertebral stripe (Supp. Fig. S5).

Thirty two transcripts were found at significantly different abundance between the *thin* and *wide* morphs (Fig. 3b; Supp. Table S2), ten of which were associated with *per3*, a gene involved in the circadian rhythm of vertebrates^30, 31^. Six transcripts were at lower concentration within the stripe compared to dorsal skin outside the stripe, and none showed a higher abundance (Fig. 3c). Three of these were transcripts of *aldoa,* a gene expressed during melanogenesis in mammals^32^ and up-regulated in yellow compared to black carp^33^.

### Recent evolution of the thin and wide alleles

To estimate the age of the *wide* and *thin* alleles and look for signatures of selection, we used Ancestral Recombination Graphs (ARG) analyses on four individuals of *P. robeensis* representative of the different phenotypes which were sequenced at a higher coverage (12.97X on average). Times to the Most Recent Common Ancestor (TMRCA) for the *thin* and *wide* alleles were more recent than the surrounding genomic regions or the total population TMRCAs in region directly surrounding the most significant SNPs outputted by the GWAS (Fig. 4a). This result is consistent with a partial selective sweep and excludes the possibility of an ancient polymorphism maintained by balancing selection. Both the *wide* and *thin* alleles have evolved recently, 100,000-300,000 years ago. These alleles thus arose long after the divergence between *Ptychadena robeensis* and its closest relatives, estimated at 3.8-8.3 million years ago^34^.

**Figure 4.**
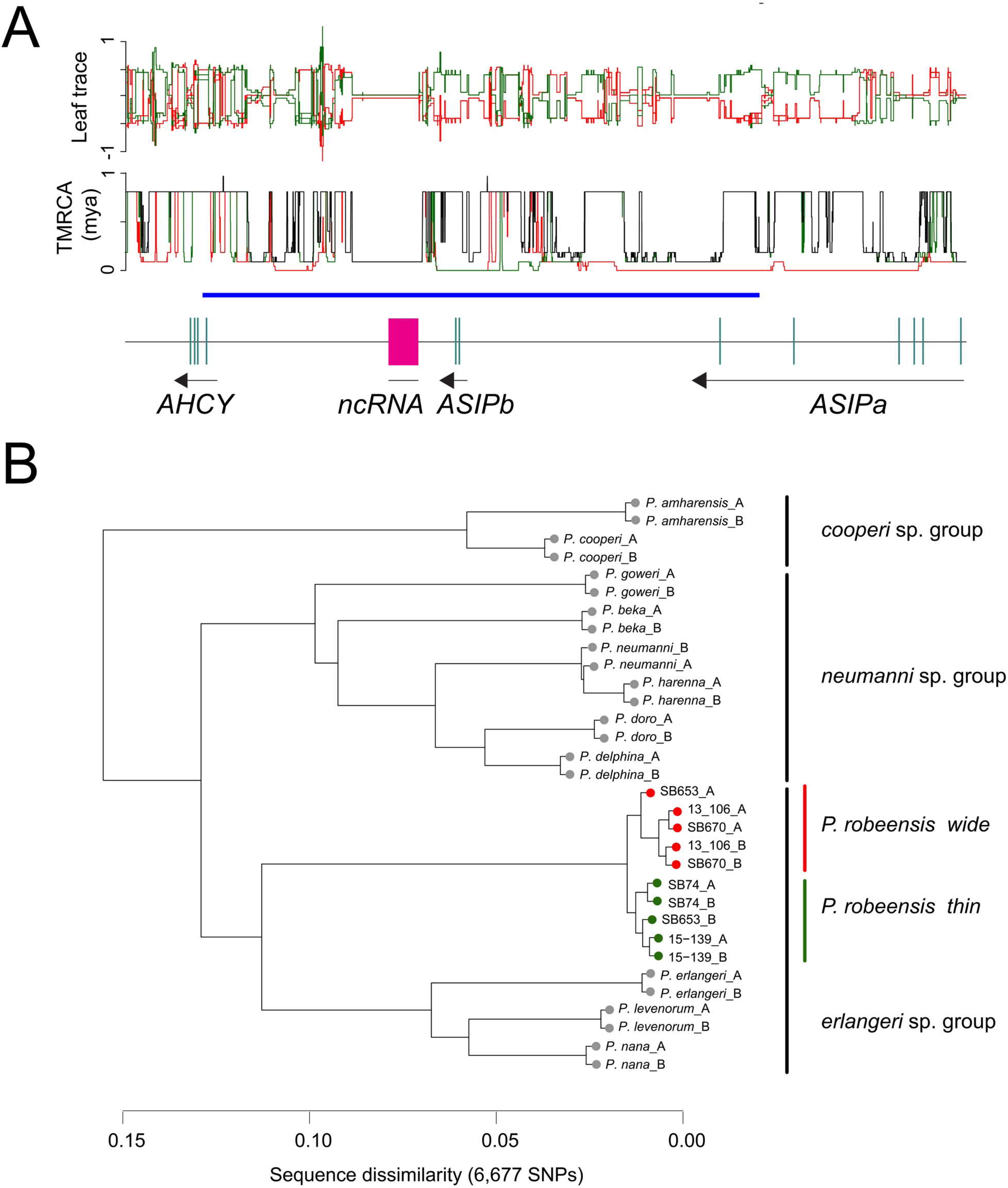
Recent evolution of the *thin* and *wide* alleles. **A** Leaf trace plot and TMRCA across the region of interest computed using ancestral recombination graphs analysis. Red and green solid lines indicate *wide* and *thin* haplotypes, respectively. Black solid lines indicate the overall population. The positions of *ASIPa, ASIPb*, *AHCY* and the ncRNA are indicated below and the region containing significant SNPs in the GWAS analysis is indicated by a horizontal blue bar. **B** Dendrograms based on dissimilarity of sequences in the region of interest (40kb region; 6,677 SNPs) in the *Ptychadena neumanni* species complex (8.29X average coverage). Haplotypes are denoted by A or B after the species or sample name. For *P. robeensis*, leaves are color-coded by haplotypes (green for *thin* and red for *wide*). Within *P. robeensis*, haplotypes are grouped by color pattern, while across the *Ptychadena neumanni* complex, they are grouped according to species relatedness and form the three clades previously described^24^ (Fig. 2a).

When comparing the genomic region of interest between all 12 species of the *P. neumanni* complex (8.29X average coverage), we found that the *thin* and *wide* alleles were indeed private to *Ptychadena robeensis* and not shared with the other *Ptychadena* species (Fig. 4.b). This result is further supported by a phylogeny of the region of interest where haplotypes are grouped according to clades within the radiation, and not phenotypes (Fig. 4.b). Additional GWAS analyses on two other members of the *P. neumanni* complex, *P. amharensis* (n = 32; 1.95X average coverage; 82,580,376 SNPs) *and P. beka* (n = 42; 4.58X average coverage; 33,430,567 SNPs), failed to detect a single locus responsible for the dorsal pattern for either species. These results indicate that the alleles found in *P. robeensis* have evolved recently and other alleles are responsible for similar dorsal patterns in closely related species.

## Discussion

In this study, we show that the anuran vertebral stripe evolved multiple times, and significantly more often in terrestrial lineages compared to terrestrial-aquatic, arboreal, and terrestrial-arboreal linages. The vertebral stripe might increase concealment from visually oriented predators, such as birds or mammals, which are more prevalent in terrestrial habitats, and thus confer a fitness advantage. The widespread polymorphism for the trait may be maintained because some predators attack preferentially the most common morph and/or because the cryptic advantage of the dorsal pattern changes across the environment or through seasons. The vertebral stripe polymorphism could also be the result of recently evolved alleles on their way to fixation. This process could be particularly slow if alleles causing the vertebral stripe are dominant^17, 21^ and the stripe adaptive advantage is only moderate. Interestingly, the vertebral stripe was lost significantly more often in arboreal lineages than in other groups, indicating a potential fitness cost of the pattern in this habitat. Other color patterns could also be selected for in arboreal habitats and the vertebral stripe could be lost as a side-effect. Although the molecular and evolutionary mechanisms may vary across lineages, a shared selective pressure favored the presence of striped morphs in terrestrial anurans.

In the grass frog *Ptychadena robeensis,* we identified two *ASIP* alleles responsible for distinct vertebral stripe morphs. In doing so, we establish the first evidence of a causal link between *ASIP* and a color pattern in amphibians. The involvement of *ASIP* in melanophore differentiation and melanin production has been extensively studied in mammals^28, 35–37^, and in rodents in particular^38–40^. Vertebral stripe pattern differentiation is likely governed by differential expression of *ASIP* as alleles differ in non-protein coding regions. However, the expression level of *ASIP* is extremely low and does not differ between morphs in adult dorsal skin. Morph-dependent differential expression of *ASIP* may thus occur at an earlier stage of the animal’s development^41^, or punctually during chromatophore differentiation resulting in an overall low level of expression in the dorsal skin. In the *Ptychadena neumanni* species complex, the vertebral stripe first appears at the final stages or after metamorphosis (personal observation based on 92 individuals identified through barcoding at different developmental stages). Investigating *ASIP’s* expression levels in the dorsal skin of metamorphic and juvenile individuals will be necessary to determine the gene’s role in the establishment of the color pattern during development.

Other genes were found to be differentially expressed (DE genes) between morphs and between skin sections (within and outside the vertebral stripe). While morph-dependent DE genes are likely to be involved in the same pathways as *ASIP*, the genes differentially expressed between sections of the dorsal skin could be produced by the different mature chromatophores. As opposed to the dorsal skin, *ASIP* expression levels were high in the ventral skin, which is, as in many other anurans, uniformly white. This dorso-ventral differential expression is comparable to expression patterns found in several species of fish which also present a white ventrum^27, 41^. Interestingly, the difference in expression level between dorsal and ventral skin was most marked for the ncRNA located downstream of *ASIPb* (Supp. Fig. S4). Together with the proximity of significant SNPs outputted by the GWAS (Fig. 3), this indicates that this ncRNA is a major actor in color pattern determination in this species, likely by having a regulatory role on *ASIPa* and/or *ASIPb*. *ASIP* thus seems to play a determining role in both the dorsal color pattern and the lack of melanization in ventral skin.

In multiple organisms, *ASIP* alleles have evolved rapidly leading to parallel evolution of similar phenotypes within species^42^ or groups of closely-related species^28^. In the *P. neumanni* species complex, species presenting the same color patterns did not share the *ASIP* alleles with *P. robeensis*. The lack of signal in the GWAS conducted for *P. amharensis* and *P. beka* could also be indicative of multiple haplotypes leading to the same phenotypes within these species, which occupies much larger distribution ranges than *P. robeensis* and presents greater intraspecific genetic variances. We estimated *P. robeensis’ ASIP* alleles to have evolved less than 300,000 years ago, much after diverging from its closest known relatives. *Ptychadena robeensis* occupies grasslands and cultivated fields where opportunities for hiding from predator under the vegetation are rare and substrate varies from grass to bare soil. A vertebral stripe (wide or thin) may thus provide a significant fitness advantage by enhancing crypsis or disrupting body shape recognition.

Recurrent evolution in the regulatory region of *ASIP* could have led to the same color patterns in this group of closely-related species, similarly to the horizontal stripes in African cichlids caused by repeated evolution at *agrp2* regulatory region^43^. However, mutations impacting the expression of genes interacting with *ASIP* (such as *MC1R* for example^27^) may also be responsible for the vertebral stripe in *Ptychadena* spp. and other terrestrial anurans. By demonstrating the involvement of *ASIP* in a widespread trait, our study opens new research avenues on color patterns in anurans. Mutations in the regulatory regions of *ASIP* or interacting genes causing the appearance or loss of the vertebral stripe are likely occurring at a high rate in anurans, making this trait an ideal system to study parallel evolution.

## Methods

### Comparative analysis of vertebral stripe evolution in anurans

We conducted a comparative analysis across the Anura using the largest dated molecular phylogeny of amphibians published to date^22^. This phylogeny comprises 3,309 species (=45% of currently recognized species^23^), representing most families, subfamilies and genera^22^. We collected data on dorsal color pattern for 2,785 of these species by examining all photographs available for each species on *Amphibiaweb* (https://amphibiaweb.org). When no or few photos were available, we searched additional sources such as original species descriptions and the number of photographs examined for each species was systematically recorded (Supp. Table S1). Dorsal color patterns were classified in the following categories: thin, medium or wide stripe, and unstriped. If a species had at least one individual counted in the *unstriped* and one of the *striped* categories, the species was considered polymorphic for the trait.

Habitat use data was collected independently for 2,620 species based on multiple large studies (Supp. Table S1). Anuran habitats were categorized as arboreal, aquatic, terrestrial, arboreal/terrestrial and terrestrial/aquatic, based on the main habitat occupied by adult individuals (when reproductive and general habitats were available).

### Ancestral state reconstruction of dorsal color patterns

To reconstruct the evolutionary history of the vertebral stripe, we created 1,000 stochastic maps of the trait onto the phylogeny using the function *make.simmap* in the R package *phytools*^44^. Because most of the *striped* species were polymorphic for the trait (78%), and as erroneous categorization was more likely for the fixed than for the polymorphic categories (if only few photos were available), we recategorized species as either having at least some individuals presenting a vertebral stripe, or without any striped individuals. We thus used two categories, *striped* (including polymorphic species) and *unstriped* with a model allowing transition rates between the two morphs to be different (ARD), and estimated the number of transitions during the evolutionary history of anurans.

### Dorsal pattern evolution in different habitats

To test the hypothesis that the dorsal stripe might be selected for in particular habitats, we first built 100 stochastic maps of habitat data for the 2,620 species categorized on the phylogeny. For each of the stochastic trees, we fitted a model for which transition rates between dorsal color patterns was independent of habitat and a model for which transition rates differed for each habitat using the *fitmultiMk* function from the R package *phytools*, and compared them using a likelihood ratio test (Supp. Figure S1). As the habitat-dependent model of evolution for the vertebral stripe was systematically better fitted than the independent model, we extracted the estimated transition rates between *striped* and *unstriped* phenotypes for each habitat and each of the 100 stochastic maps. Because few species were categorized as aquatic (1.87 % of included taxa, i.e. 49 species), the variance in transition rate estimates was much greater than for the other habitats (Supp. Figure S2), so we excluded aquatic lineages from further analyses. We compared the transition rates between pairs of habitats using a Tukey honest significant differences test (Table 1).

### Sampling of Ethiopian Ptychadena

Individuals of the *Ptychadena neumanni* species complex were collected in the highlands of Ethiopia between 2011 and 2019. Our study was approved by the relevant Institutional Animal Care and Use Committee at Queens College and New York University School of Medicine (IACUC; Animal Welfare Assurance Number A32721–01 and laboratory animal protocol 19–0003). Frogs were sampled according to permits DA31/305/05, DA5/442/13, DA31/454/07, DA31/192/2010, DA31/230/2010, DA31/7/2011 and DA31/02/11 provided by the Ethiopian Wildlife Conservation Authority. We photographed individuals in life and euthanized them by ventral application of 20% benzocaine gel. We extracted tissue samples and stored them in RNAlater or 95% ethanol. Adult individuals were fixed in 10% formalin for 24 to 48 hours, and then transferred to 70% ethanol. After preservation, we took additional photographs of all individuals. All specimens were deposited at the Natural History Collection of the University of Addis Ababa, Ethiopia. Tissue samples are deposited at the Vertebrate Tissue Collection, New York University Abu Dhabi (NYUAD).

### Histological skin sections

Dorsal and ventral skin sections were extracted from ten preserved adult specimens: two thin striped (SB81, SB82) and one unstriped (SB69) *Ptychadena robeensis* specimens, two thin striped (SB493, SB510) and one wide striped (SB494) *P. nana*, two wide striped (SB552, SB548) *P. erlangeri*, and two wide striped (SB584, SB593) *P. amharensis.* The skin samples were embedded in paraffin blocks and sections of 4 µm thickness were produced. The sections were stained with hematoxylin-eosin (HE) and chromatophores were examined using a Leica DMI 6000 B microscope.

### DNA and RNA extractions and sequencing

Genomic DNA of 61 *Ptychadena robeensis* individuals was extracted from liver tissue using the DNeasy blood and tissue kit (Qiagen, Valencia, CA). RNA was extracted from the skin of 13 individuals using a RNeasy mini kit (Qiagen, Valencia, CA). For eight individuals, RNA was extracted within and outside the vertebral stripe separately, for three individuals lacking any dorsal pattern, a single sample of dorsal skin was used, and for two individuals, RNA from ventral skin was extracted. We quantified extracted DNA and RNA using a Qubit fluorometer (Life Technologies). Libraries were prepared using a NEB library prep kit and sequenced on Illumina NextSeq 550 flow cells at the Genome Core Facility of New York University Abu Dhabi. After quality filtering, reads were aligned to the *de novo* assembly *Ptychadena robeensis* reference genome^25^. The average coverage of the genomic data was 4.84X, except for four individuals which we sequenced at an average of 12.97X. Variants were called using the function *HaplotypeCaller* from *gatk v3.5*^45^. The low-coverage and higher-coverage samples were then combined and genotyped in two separate datasets using *CombineGVCF* and *GenotypeGVCFs* functions from *gatk*.

### Genome-wide association study on P. robeensis

After examination of the low-coverage genomic dataset (n=61 individuals; SNPs PCA and visual examination in IGV), we realized that five individuals were hybrids, likely resulting from the crossing of *P. robeensis* and *P. levenorum*, a closely related species with a partially overlapping distribution range^24^. After removing the hybrids, the dataset comprised 17,797,568 SNPs. We excluded four individuals, which presented no dorsal melanization and could not be categorized as thin or wide striped, and our final dataset contained 52 individuals (25 wide striped and 27 thin striped. Stripe phenotype (thin, wide or unstriped) and coloration (brown or green) were not correlated. Quality checks and the genome-wide association study (GWAS) were done using *PLINK 1.9*^40, 41^. We checked for individual relatedness as well as major discrepancies between samples in data missingness and minor allele frequency. However, because of the low-coverage nature of our data, we did not apply any stringent quality filtering. We extracted and visualized the result of the GWAS using the R package *qqman*^48^.

### Test for selection and alleles ages in Ptychadena robeensis

We searched for signatures of selection in the regions linked to dorsal stripe pattern and determined the age of the alleles using *ARGweaver*. In short, *ARGweaver* reconstructs a set of Ancestral Recombination Graphs (ARGs) for every non-recombining interval in the genomic region of interest. The program then samples from the posterior distribution of ARGs given an evolutionary model. Regions under positive selection should display a reduced coalescence time whereas regions under ancient balancing selection should have older coalescence time compared to neutral regions.

We ran *ARGweaver* on the high-coverage *Ptychadena robeensis* dataset (12.97X average coverage) comprising four individuals. We used the mutation rate estimated for the species group^34^ 6.98e-10/bp/generation (with a 2-year generation time) and the average recombination rate estimated for *Xenopus tropicalis*^49^, 9.73e-9/bp/generation as priors. The effective population size was estimated as a function of time in SMC++^50^ elsewhere^51^, and we used a maximum coalescence time of 4 million generations, around twenty times the harmonic mean of the estimated effective population size over time. *ARGweaver* was run for 5,000 iterations and ARGs were sampled every 100 iteration. Convergence of the chain was monitored visually by plotting multiple ARG statistics (priors, likelihood, number of recombination events, total branch length, number of variant sites not explained by a single mutation) against iteration number. All statistics were stationary after the first 2,000 iterations, so we discarded these 2,000 first iterations as burn-in for further analyses.

We inspected the phased haplotypes and categorized them in *wide* or *thin* haplotypes, thereby considering haplotypes rather than individuals in the subsequent analyses. We extracted times to the most recent common ancestor (TMRCA) for the *wide* and *thin* haplotypes, as well as for the whole population and compared the three curves across our region of interest. Ancient balancing selection should show TMRCA older than neutral genomic regions and an equal TMRCA between the haplotypes and the overall population. Positive selection or recent balancing selection, on the other hand, should have and overall TMRCA similar to neutral regions but haplotype TMRCAs more recent than the overall TMRCA.

### Phylogeny of haplotypes

In order to compare the region of interest across the *Ptychadena neumanni* species complex, we reconstructed a haplotype phylogeny. The genomes of the 12 species (one individual per species, except for *P. robeensis* for which the genomes of five individuals were included) were phased using Beagle 5.1^52^. We then built a phylogeny of haplotypes in the region of interest (40kb region; 6,677 SNPs) using the R package *SNPRelate*^53^ (Fig. 4b).

### Gene expression analysis

Reads were aligned to the annotated reference genome using HISAT2^54^ and StringTie2^55^. A transcriptome-wide gene count matrix was then created using the script *prepDE.py3* provided on the StringTie website. Subsequent analyses were conducted in the *R* environment^56^. We used the *R* package *edgeR*^57^ to filter and normalized our data prior analysis. We filtered out genes which had a count inferior to 1 count-per-million (cpm) in at least 16 samples (>50% of the 21 samples in total) and applied a “Trimmed Mean of M-values” (TMM) normalization of the data using the *R* package *DESeq2*^58^. We then identified differentially expressed (DE) genes between wide and thin striped individuals as well as between sections of dorsal skin within and outside the vertebral stripe using the *DESeq* function of the same package.

### Annotation of the region of interest

In order to determine the type of variant responsible for the vertebral stripe pattern in *Ptychadena robeensis*, we annotated the region of chromosome 11 containing significant SNPs in the GWAS. We visually examined the transcripts obtained from our RNAseq data against the annotation for protein coding sequences predicted by Augustus^25, 59^ in the region using IGV^60^. Significant SNPs outputted by the GWAS were all located outside protein-coding regions, in between three genes: two predicted genes 40 kb apart were identified as *ASIP* and a third, 38kb downstream, was identified as *AHCY* using MegaBlast^61^.

While a single *ASIP* gene is known in birds and mammals, two genes, *ASIP1* and *ASIP2* are present in teleost fish^27, 62^. In *Xenopus tropicalis*, a single *ASIP* gene is predicted, but no focal study in amphibians has been conducted to date. In order to determine whether the two genes in *P. robeensis* correspond to *ASIP*, *ASIP1,* or *ASIP2*, we translated the genomic sequences and produced a protein alignment and maximum likelihood phylogeny with the *ASIP* gene family in vertebrates using *seaview*^63^ (Supp. Fig. S3). Both genes grouped together and with other amphibians’ *ASIP*. We thus named them *ASIPa* and *ASIPb* to avoid any confusion with *ASIP1* and *ASIP2* from teleost fish. *ASIPa* is composed of six exons, two of which are copies of *ASIPb*’s only two exons. Additionally, we detected a non-coding RNA (ncRNA) downstream *ASIPb*. A protein alignment revealed that this ncRNA contains a region aligning with the third exon of *ASIP* in other vertebrates, which is absent from *ASIPa* and *ASIPb*.

### Quantitative real-time PCR experiment

Quantitative real time PCR (qPCR) was conducted on RNA extracted from dorsal skin within (n=2) and outside (n=4) the vertebral stripe, and ventral skin (n=1). Each reaction was triplicated to minimize the impact of experimental error. Two reference genes were selected using our RNAseq dataset with the following criteria: a minimum of 50 count per million in all samples, the lowest possible variance in expression level among samples and a minimum of two exons. Candidate reference genes were then checked for functional independence and compared to genes typically used for qPCR in *Xenopus laevis*. We retained *rpl27* and *abce1* as candidate reference genes.

Primers for *ASIPb, rpl27* and *abce1* were designed based on the *Ptychadena robeensis* reference genome and annotation using Primer3Plus^64^. Primers were designed to span an exon-intron junction to avoid amplification of genomic DNA during the qPCR. The experiment was run using a StepOnePlus real-time PCR system and a Power SYBR Green RNA-to-CT 1-step kit (Applied Biosystems) on a volume of 20µl. Results were analyzed using the R package *pcr*^65^. We compared the expression levels of *rpl27* and *abce1* across samples and retained *rpl27*. Relative expression levels of *ASIPb* between our samples were calculated using *rpl27* as reference gene and dorsal skin outside the stripe as reference group as it should have the lowest level of *ASIPb*.

## Acknowledgments

We would like to thank the Ethiopian Wildlife Conservation Authority and the Ethiopian Biodiversity Institute for providing us with collecting and export permits for the samples. Fieldwork in Ethiopia would not have been possible if not for the invaluable assistance of Megersa Kelbessa, Itbarek, and Samuel Woldeyes of Rock Hewn Tours. We also thank the important number of students and postdocs who collected *Ptychadena* specimens and samples over the years, and in particular Xenia Freilich, Jacobo Reyes-Velasco, Justin Wilcox, Sebastian Kirchhof and Marcin Falis. We are very thankful for the help from Marc Arnoux and Nizar Drou, from the Genome Core Facility and the Bioinformatics group at NYUAD. This research was carried out on the High-Performance Computing resources at New York University Abu Dhabi. We also thank David Howse and Sayel Daoud of The National Reference Laboratory and Rachid Rezgui from the Microscopy Core Facility at NYUAD for their help in producing and visualizing histological sections.

## Author contributions

SG and SB designed the study. KDU and SG collected color pattern and habitat data. SG, YB, and SB collected *Ptychadena* spp. specimens and samples. SG extracted DNA and RNA from *Ptychadena* spp. samples and ran the and qPCR. SG produced and interpreted histological section photographs and conducted comparative, genomic and gene expression analyses. IH and YB provided help and advice on genomic and transcriptomic analyses. All authors read and contributed to the manuscript.

## Supplemental information

**Table S1. Phenotype and habitat data and references.**

**Figure S1.**
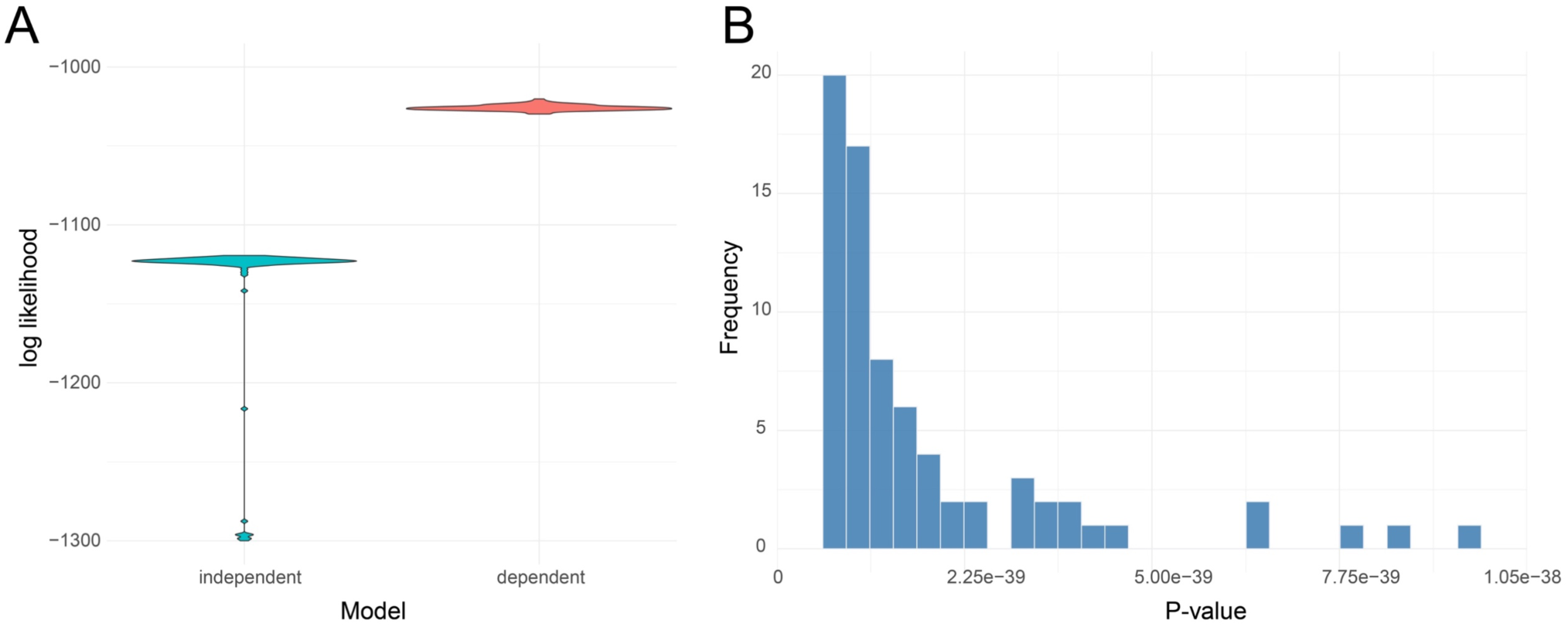
Gain and loss rates of the anuran vertebral stripe are habitat-dependent. **A** Likelihood of the habitat-dependent and independent models estimated for 100 stochastic maps. **B** P-values of the 100 likelihood ratio tests between dependent and independent. For all 100 stochastic maps, the model with transition rates dependent of the habitat is favored.

**Figure S2.**
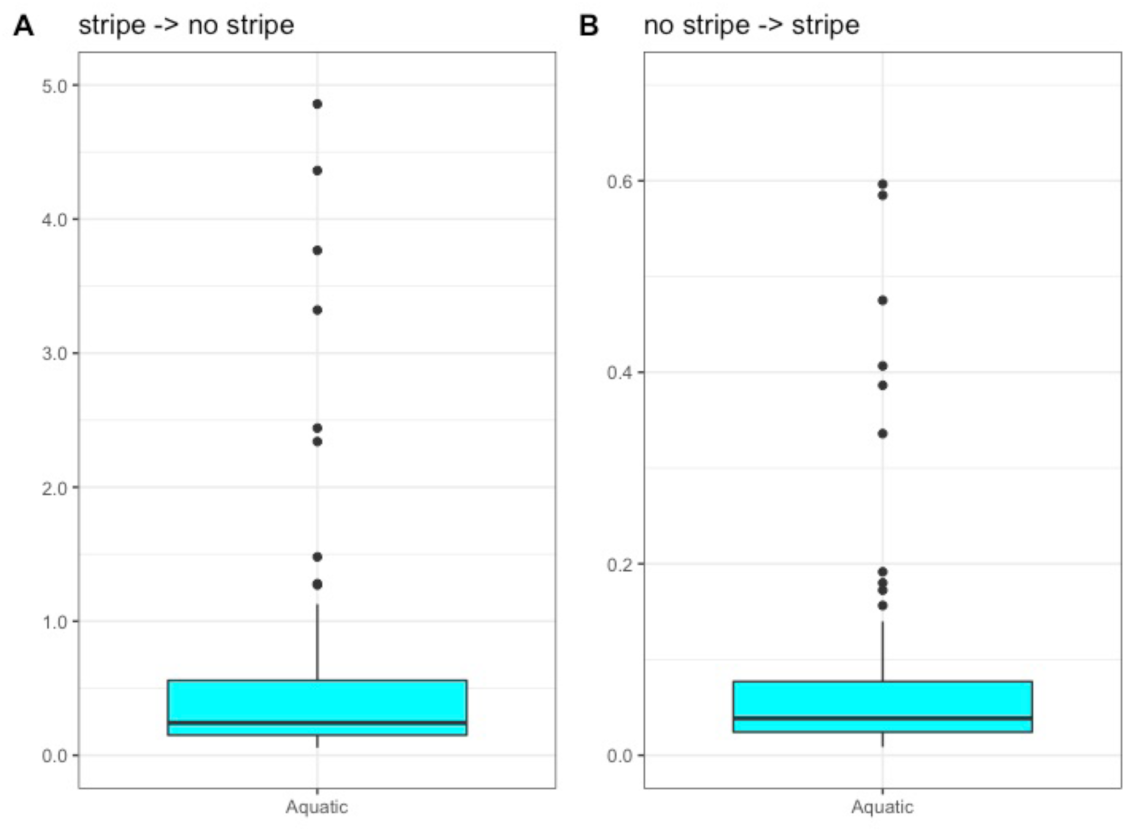
Transition rates between striped and unstriped phenotypes for aquatic lineages estimated for 100 stochastic maps. The low number of species in the aquatic category (1.87 % of included taxa, i.e. 49 species) results in important variation in rates among the stochastic maps.

**Figure S3.**
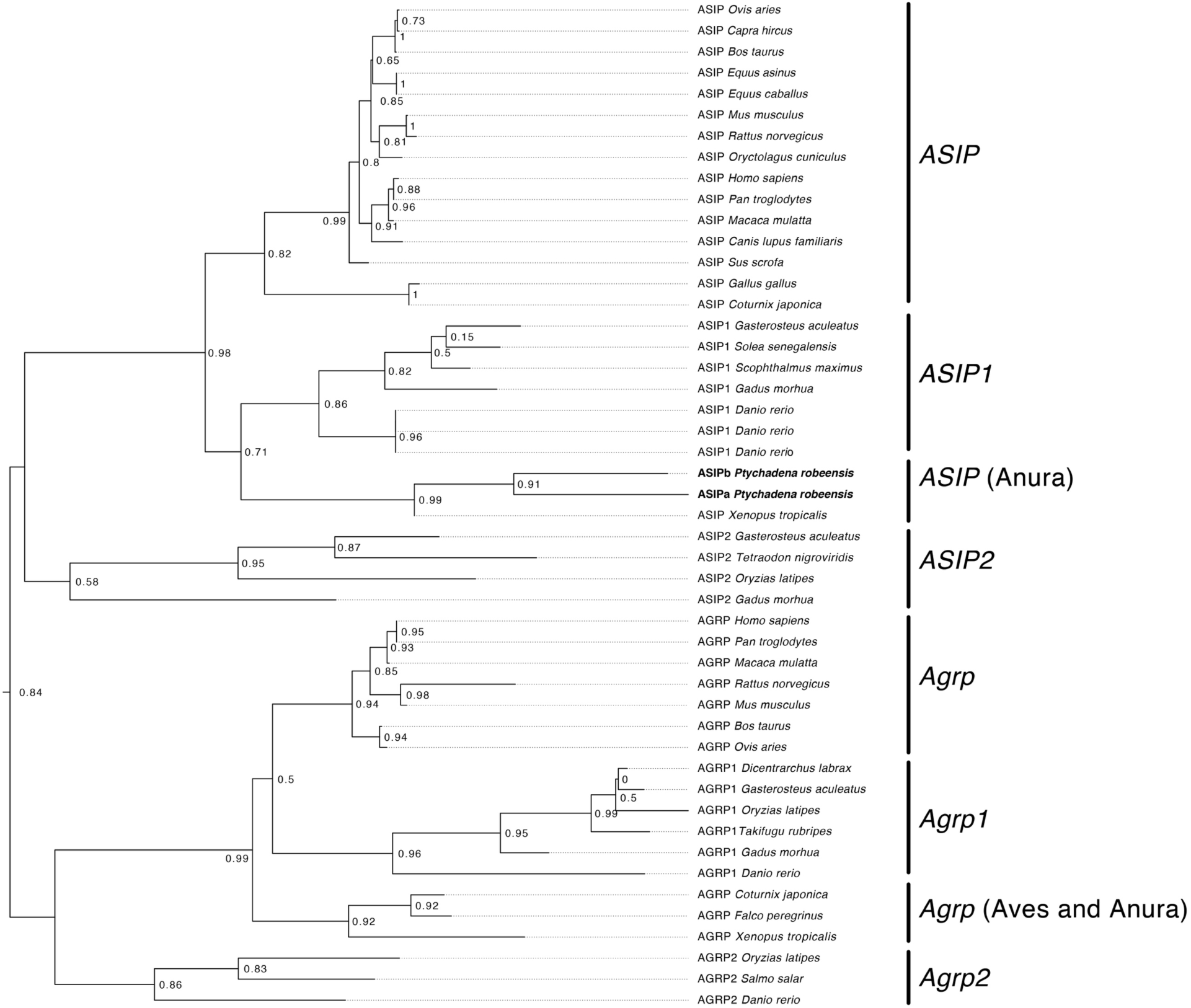
Position of *ASIPa*and *ASIPb* in the evolution of *ASIP.* Maximum likelihood tree based on a protein alignment using MUSCLE^66^. *ASIPa* and *ASIPb* are grouped with *ASIP* of *Xenopus tropicalis,* excluding the possibility for the two genes to correspond to fish’s *ASIP1* and *ASIP2*. Examination of the protein alignment showed that *ASIPa* and *ASIPb* share two exons, resulting from a gene duplication event.

**Figure S4.**
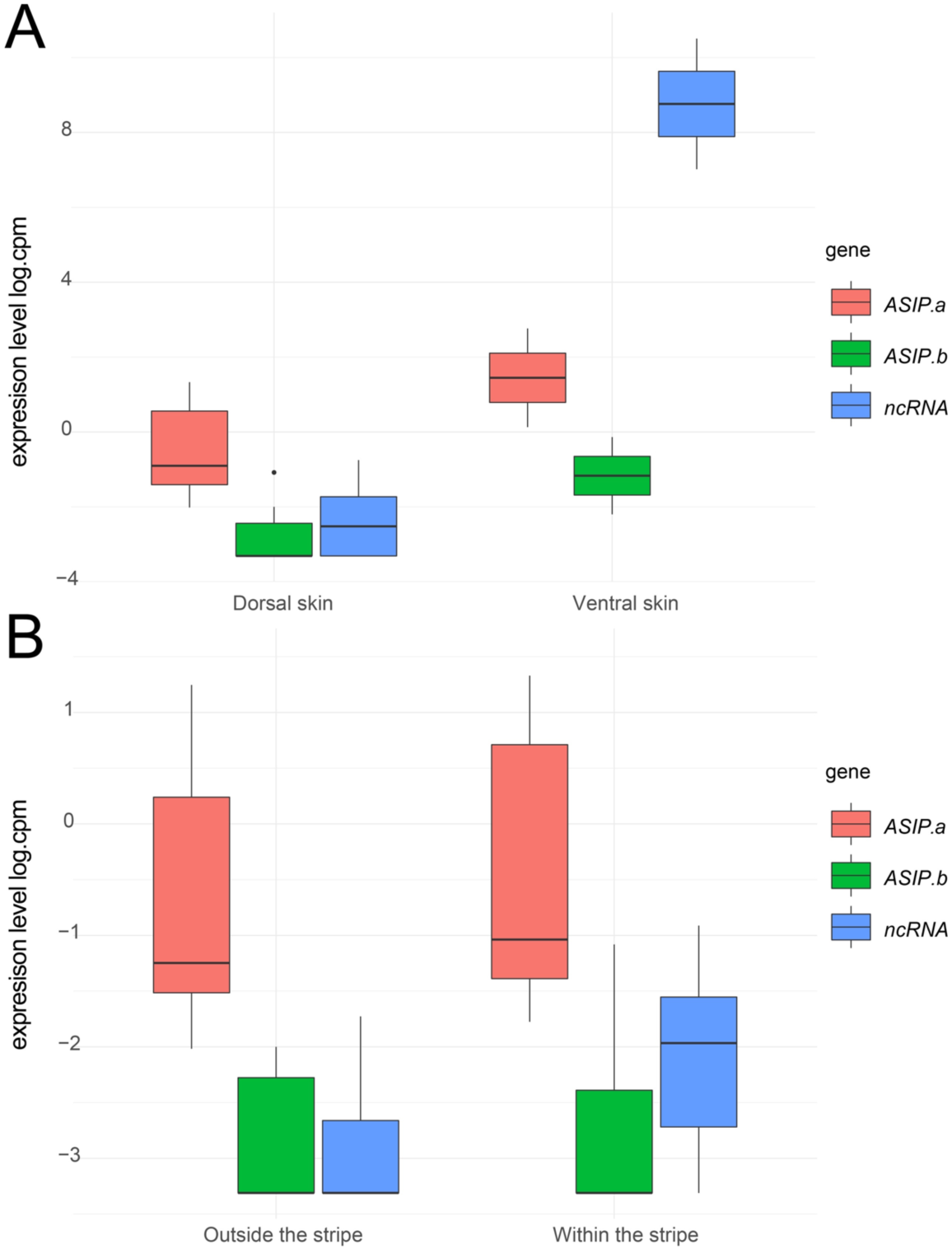
Expression levels of ASIP in the skin of *Ptychadena robeensis*. Normalized expression levels of *ASIPa*, *ASIPb* and the ncRNA from RNAseq data is given in log count per million. **A** ventral skin (n=2) shows a greater number of all three transcripts than dorsal skin (n=19), **B** skin within (n=4) and outside (n=4) the vertebral stripe do not show significant differences in expression level of *ASIPa*, *ASIPb* or the ncRNA.

**Figure S5.**
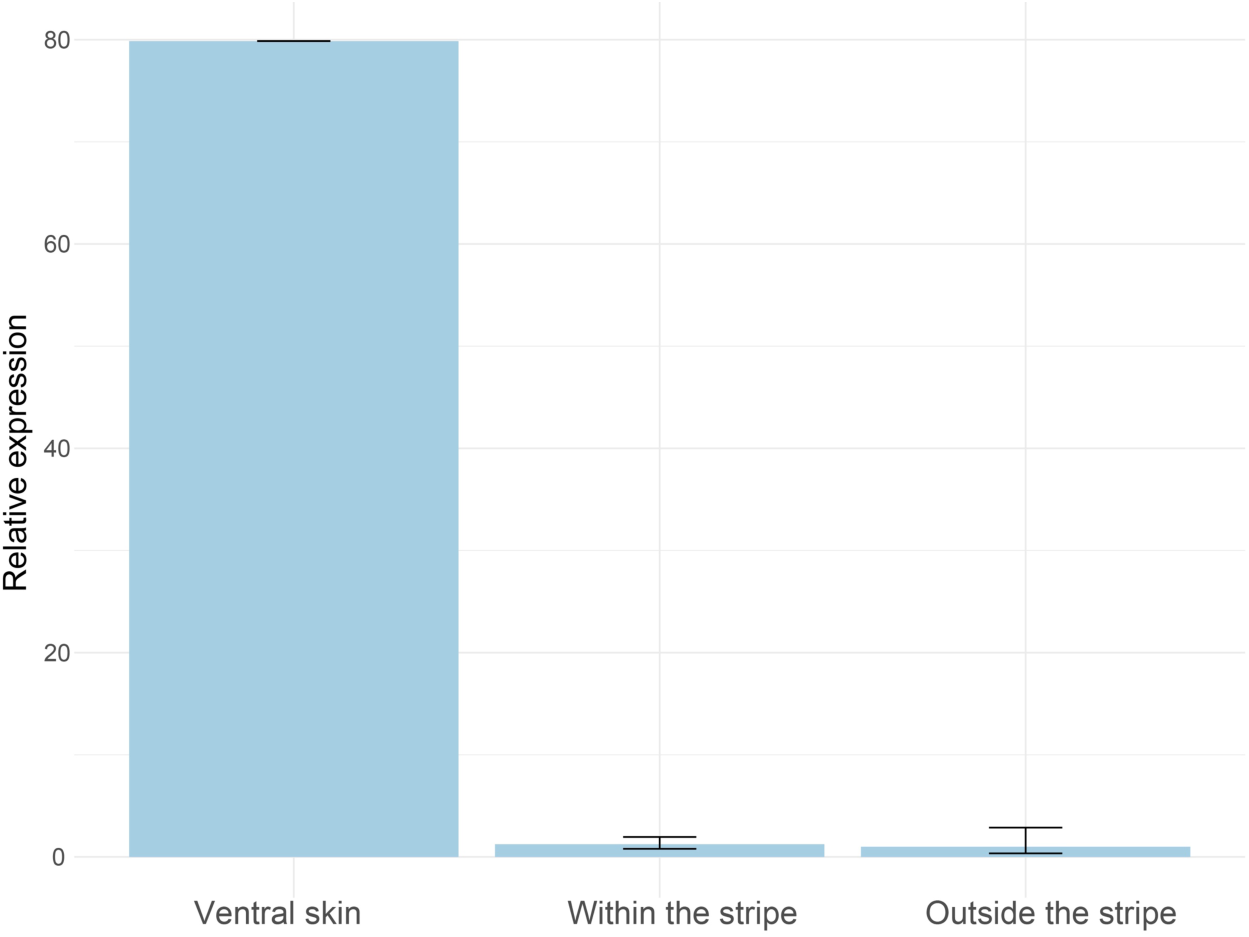
Quantitative real-time PCR of *ASIPb* in *Ptychadena robeensis*. Relative expression levels of *ASIPb* in ventral skin (n=1) and dorsal skin within (n=2) and outside (n=4) the vertebral stripe measured by qPCR. Each reaction was triplicated and average CT value for each individual was used. *ASIPb* expression level in ventral skin is 80 times greater than in dorsal skin, while no significant differential expression is detected within versus outside the vertebral stripe.

